# SARS-CoV-2 ribosomal frameshifting pseudoknot: Improved secondary structure prediction and detection of inter-viral structural similarity

**DOI:** 10.1101/2020.09.15.298604

**Authors:** Luke Trinity, Lance Lansing, Hosna Jabbari, Ulrike Stege

## Abstract

Severe acute respiratory syndrome coronavirus 2 (SARS-CoV-2) has led to the COVID-19 pandemic; a pandemic of a scale that has not been seen in the modern era. Despite over 29 million reported cases and over 900, 000 deaths worldwide as of September 2020, herd immunity and widespread vaccination efforts by many experts are expected to be insufficient in addressing this crisis for the foreseeable future. Thus, there is an urgent need for treatments that can lessen the effects of SARS-CoV-2 in patients who become seriously affected. Many viruses including HIV, the common cold, SARS-CoV and SARS-CoV-2 use a unique mechanism known as −1 programmed ribosomal frameshifting (−1 PRF) to successfully replicate and infect cells in the human host. SARS-CoV (the coronavirus responsible for SARS) and SARS-CoV-2 possess a unique RNA structure, a three-stemmed pseudoknot, that stimulates −1 PRF. Recent experiments identified that small molecules can be introduced as antiviral agents to bind with the pseudoknot and disrupt its stimulation of −1 PRF. If successfully developed, small molecule therapy that targets −1 PRF in SARS-CoV-2 is an excellent strategy to improve patients’ prognoses. Crucial to developing these successful therapies is modeling the structure of the SARS-CoV-2 −1 PRF pseudoknot. Following a structural alignment approach, we identify similarities in the −1 PRF pseudoknots of the novel coronavirus SARS-CoV-2, the original SARS-CoV, as well as a third coronavirus: MERS-CoV, the coronavirus responsible for Middle East Respiratory Syndrome (MERS). In addition, we provide a better understanding of the SARS-CoV-2 −1 PRF pseudoknot by comprehensively investigating the structural landscape using a hierarchical folding approach. Since understanding the impact of mutations is vital to long-term success of treatments that are based on predicted RNA functional structures, we provide insight on SARS-CoV-2 −1 PRF pseudoknot sequence mutations and their effect on the resulting structure and its function.

## 1 Introduction

The COVID-19 pandemic caused by the novel severe acute respiratory syndrome coronavirus 2 (SARS-CoV-2) has created a health emergency. Patients who are affected by COVID-19 can develop symptoms including severe pneumonia, acute respiratory distress syndrome, or suffer multiple organ failure [1, 2, 3]. Currently, there is no specific treatment for COVID-19 [4]. In order to decrease the mortality of patients who are seriously affected, there is an urgent need to develop SARS-CoV-2 antiviral drug therapies. Sequence analysis of the SARS-CoV-2 genome classify it as a member of the Betacoronavirus subfamily, which includes SARS-CoV and MERS-CoV [5, 6]. Despite having one of the lowest fatality rates among Betacoronaviruses (MERS fatality rate 34.3% [7], SARS fatality rate 11% [8], COVID-19 fatality rate, approximately 2.3% [9]), SARS-CoV-2 caused a pandemic because it is highly contagious. The *R*_0_ *basic reproduction number* (number of cases resulting from each index case) for COVID-19 is between 2 and 2.5 compared with 0.7 for MERS [6]. This explains why there have been less than 3000 cases of MERS [6], and over 29 million COVID-19 cases worldwide. Despite their differences in mortality and transmissibility, all coronaviruses use a replication strategy that is a promising target for therapeutic drug development [10].

One key step in the replication of coronaviruses is *ribosomal frameshifting*. Ribosomes are macromolecules of RNAs and proteins found in abundance in living cells. The function of ribosomes is to translate RNAs into proteins during a process called translation. During translation, there is a ribosome elongation cycle that occurs in which the ribosome translates sets of three nucleotides at a time into their corresponding amino acids. Each set of three nucleotides is referred to as a codon. Translation begins when a ribosome binds to a start codon. Then, each codon is translated in turn by the ribosome creating a chain of amino acids known as a polypeptide. The elongation cycle ends when the ribosome reaches a stop codon. The sections of consecutive codons that have the ability to be translated are known as open reading frames (ORF). An *open reading frame* is a continuous stretch of codons that begins with a start codon and ends at a stop codon.

In the case of the SARS-CoV-2 there are two different long open reading frames comprising two-thirds of the viral genome: ORF1a and ORF1b (see Figure 1) [11]. ORF1b is out of frame with respect to ORF1a, meaning the ribosome will not translate both frames without shifting from one frame to the other. All coronaviruses utilize the combination of a slippery sequence (the sequence where the ribosome is prone to slipping forwards or backwards into a different reading frame) and an *RNA pseudoknot* to cause the ribosome to shift from one reading frame into the other [12, 13]. Usually during translation, the ribosome unwinds the pseudoknot structure and continues translation without shifting reading frames. Sometimes, the pseudoknot causes the ribosome to pause at the slippery sequence, and then shift backwards one nucleotide (referred to as ‘−1’) before continuing in the second reading frame. This molecular mechanism is called *ribosomal frameshifting*. This shift results in the creation of a fusion ORF1a/1b polypeptide that is vital for viral replication [14]. The site where this −1 programmed ribosomal frameshift (−1 PRF) occurs has great promise as a therapeutic target, as decreasing the rate at which the ribosomal frameshift occurs, referred to as the −1 PRF efficiency, can inhibit viral replication [15, 16].

**Figure 1:**
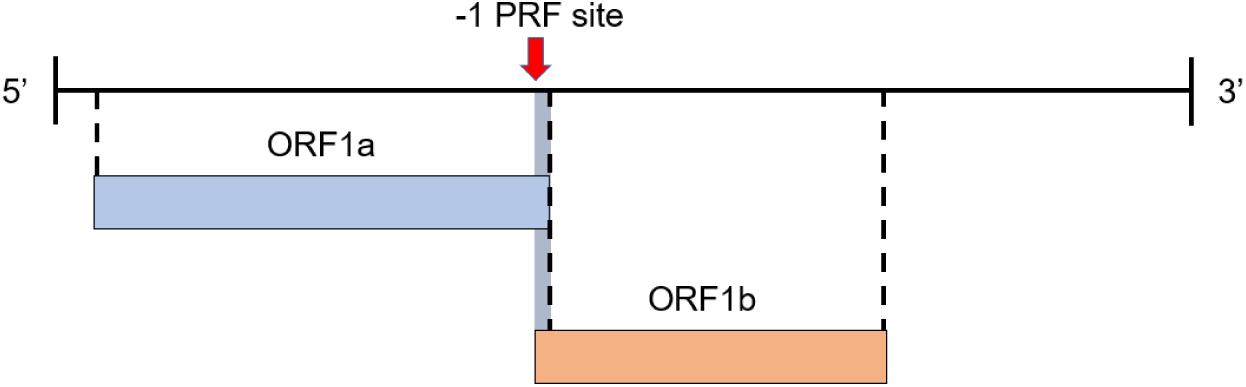
SARS-CoV-2 Viral Genome. The −1 programmed ribosomal frameshifting (−1 PRF) site is marked by a red arrow. Normally, the ribosome translates the complete ORF1a. Sometimes, at the −1 PRF site, the ribosome may shift from ORF1a to ORF1b. This shift results in the synthesis of a fusion ORF1a/1b polypeptide.

The pseudoknot that causes ribosomal frameshifting is an example of an RNA *secondary structure*. The primary structure of RNA is a sequence of nucleotide bases connected to one another through the sugar backbone. When RNA folds back on itself, complementary bases pair, resulting in formation of the secondary structure. The most common RNA secondary structure is the hairpin loop. A *hairpin loop* occurs when RNA folds to form a double helix called a stem, with the unpaired nucleotides forming a single stranded region called a loop (see Figure 2a). RNA secondary structures that are crucial to inducing −1 PRF are most commonly identified as a specific type of pseudoknot referred to as *H-type*. The pairing of unpaired bases of a hairpin loop with complementary bases downstream, results in an H-type pseudoknot (see Figure 2).

**Figure 2:**
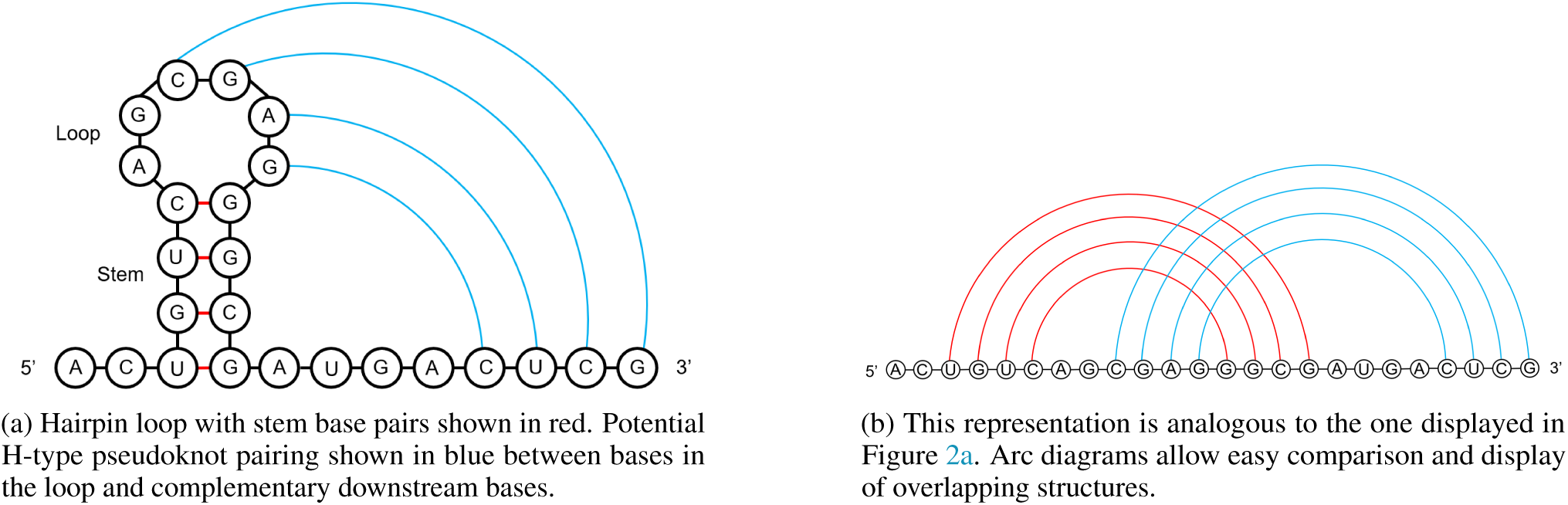
H-type pseudoknot forming from hairpin loop (a) and represented in conventional arc diagram format (b).

Pseudoknotted base pairs are referred to as crossing base pairs, because when viewed in a conventional arc diagram, the bases cross one another. As the ribosome moves along the RNA during elongation, the pseudoknotted structure represents a kinetic barrier and must be disrupted to allow the next codon into the decoding center, i.e., *translocation* [17].

RNA folding refers to the process by which RNA acquires secondary structure through interacting nucleotide bases. RNA folding can be described sequentially, with the *initial* stem forming first (Figure 2 red lines) followed by an additional stem (Figure 2 blue lines). This concept is supported based on the *hierarchical folding hypothesis*. This hypothesis posits that RNA first folds into a pseudoknot-free structure, one with no crossing base pairs (e.g. hairpin loop). Then, additional bases pair and may form pseudoknots to lower the minimum free energy of the structure [18]. This hypothesis is supported by experts [19, 20, 21], as well as by experiments on an H-type frameshifting pseudoknot where it was observed that the initial stem formed first followed by the second pseudoknotted stem [22].

The expected secondary structures of −1 PRF stimulating pseudoknots, referred to as native or wild-type pseudoknots, have been identified and studied for a wide variety of viruses [23, 24]. In addition, it has been identified that some non-native pseudoknot conformations, those that differ from the native structure, play a role in regulating frameshifting. Specifically, the conformational plasticity of pseudoknots, i.e., their ability to fold into non-native structures, was found to be correlated with −1 PRF efficiency [23]. Potential non-native secondary structures for the SARS-CoV-2 −1 PRF pseudoknot have yet to be clearly identified.

The latest research on the SARS-CoV-2 −1 PRF pseudoknot utilized 3-D physics-based modeling to identify three unique stable structural conformations [25]. However, these results do not account for non-native pseudoknot conformations in the initial secondary structure landscape. Specifically, Omar et al. in 2020 used the native secondary structure of the SARS-CoV-2 −1 PRF pseudoknot as a starting point in their structure prediction [25]. Given that non-native folding pathways correlate with −1 PRF efficiency, a devoted study into non-native pseudoknot conformations is valuable in structural prediction. However, as of yet there has not been a dedicated presentation of non-native pseudoknot conformations for the SARS-CoV-2 −1 PRF pseudoknot.

In addition, mutations have been reported in all components of the SARS-CoV-2 −1 PRF pseudoknot [26, 27]. Yet, the possible effects of such mutations on the RNA structure formation and resulting secondary structures have not been fully investigated. Understanding the impact of mutations is vital to long-term success of treatments that are based on predicted RNA structures. Interestingly, the most prevalent mutation to the SARS-CoV-2 −1 PRF pseudoknot sequence was found to increase the similarity of the SARS-CoV-2 −1 PRF pseudoknot structure to that of the MERS-CoV −1 PRF pseudoknot structure [26]. Further validation of the similarity between the SARS-CoV-2 and MERS-CoV −1 PRF pseudoknots is highly relevant for future coronavirus structure prediction efforts.

Previous work found that reducing the frameshifting efficiency of the original SARS-CoV attenuated viral propagation to a significant degree [28, 15, 16]. Small molecule therapy has been shown to be effective for SARS-CoV-2 in a lab setting, where introducing a specific molecule to limit the conformational plasticity of the pseudoknot reduced −1 PRF by 60% [29]. In addition, the small molecule binding was found to be resilient to mutations in the SARS-CoV-2 −1 PRF sequence [27].

These experimental results demonstrate the potential for SARS-CoV-2 treatments based on −1 PRF inhibition. Increasing comprehensive knowledge of predicted SARS-CoV-2 −1 PRF pseudoknot structures will be valuable in validating existing and future small molecule therapeutics.

In this paper we identify structural similarities between the −1 PRF pseudoknots of SARS, SARS-CoV-2, and MERS-CoV. Following the hierarchical folding hypothesis (see Materials and Methods), we present structures for SARS-CoV-2 and MERS-CoV −1 PRF pseudoknots (see Results). In addition, we provide pseudoknot predictions for six SARS-CoV-2 −1 PRF mutated pseudoknot sequences (see Results). These mutations were selected because they have been observed in the population as well as experimentally validated for their effect on −1 PRF frameshifting [27]. Finally, we discuss the relationship between each set of predicted initial stems and pseudoknotted structures (see Discussion).

In order to follow up on the hypothesized link between the SARS-CoV-2 and MERS-CoV −1 PRF pseudoknot structures [26], we enumerate the specific loci of similarity as well as a consensus structure. Where previous work has identified the importance of non-native pseudoknotted structures, our research goes further by explicitly identifying non-native structures and predicting how the structure is realized. Our results contribute to RNA structural prediction by providing a complete landscape of notable but previously unidentified non-native secondary structures, that may play a role in regulating frameshifting [23]. Our pseudoknot structural predictions provide alternate starting points and input constraints which can improve the accuracy of existing 3-D physics-based modeling software packages [30, 31, 32, 33, 34, 35]. Identifying potential initial stems and non-native conformations of the SARS-CoV-2 −1 PRF pseudoknot, while accounting for prevalent mutations to the SARS-CoV-2 virus, will increase the accuracy of RNA structure prediction. Our specific contributions are as follows:

1. Structural similarity and consensus structure identification for the SARS-CoV, SARS-CoV-2, and MERS-CoV −1 PRF pseudoknots.
2. Secondary structure prediction for the SARS-CoV-2 and MERS-CoV −1 PRF pseudoknots following the hierarchical folding hypothesis.
3. Secondary structure predictions for six SARS-CoV-2 −1 PRF pseudoknot mutated sequences.
4. List of relationships between initial stems and secondary structure conformations of the SARS-CoV-2 and MERS-CoV −1 PRF pseudoknots.

We first provide appropriate background information for our secondary structure analysis of the −1 PRF pseudoknots. We then define our methodology for identification of structural similarity between SARS-CoV, SARS-CoV-2, and MERS-CoV. Next, we define our hierarchical approach in identifying the initial stems and using this information to predict secondary structure. Our first result is the region of structural similarity between the coronaviruses and their consensus structure. The second section of our results consists of predicted initial stems and secondary structures for the SARS-CoV-2 and MERS-CoV −1 PRF pseudoknots. Finally, we discuss our findings as they relate to the existing RNA secondary structure literature.

## 1.1 Background

At first it was believed that mechanical stability of frameshift stimulating pseudoknots, how much force they can withstand before unfolding, may be related to the rate at which they induce −1 PRF [36]. Experimental data disproved this hypothesis identifying that the conformational plasticity of pseudoknots, i.e., their ability to fold into non-native structures, was correlated with −1 PRF efficiency [23]. One explanation is that ribosomal complex state transitions are modulated by pseudoknot unwinding [24]. It is predicted that pseudoknots with higher plasticity are more likely to fold into alternative stable structures that can produce a higher energy barrier in late stage translocation, which in turn initiates translation in the −1 frame [37]. This aligns well with the previously held hypothesis that the unwinding pseudoknots may regulate −1 PRF by changing the tension between RNA and ribosomes [38, 39]. The formation of triple-stranded helices has been proposed to contribute to conformational plasticity by enhancing the unwinding resistance of a pseudoknot to the translocating ribosome, contributing to conformational rearrangement [22].

The native structure of the frameshift stimulating pseudoknot in SARS-CoV and SARS-CoV-2 possesses a unique three-stemmed structure (see Figure 3). The SARS-CoV −1 PRF pseudoknot sequence and native structure is conserved in SARS-CoV-2 with only a single base change in the 68 nucleotide sequence [29]. We leverage this structural conservation and the corresponding literature to conduct analysis of the SARS-CoV-2 −1 PRF pseudoknot that can contribute to treatments inhibiting viral replication.

**Figure 3:**
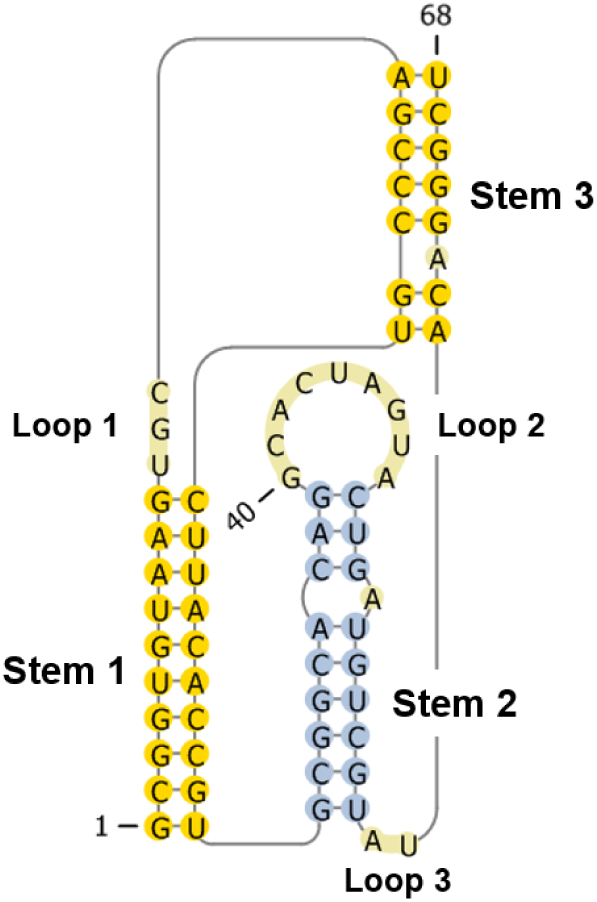
Native structure of the SARS-CoV-2 −1 PRF stimulating pseudoknot. Visualization generated using Pseu-doViewer3 [40].

Multiple small molecules have been identified that can bind with the SARS-CoV −1 PRF pseudoknot and disrupt its function [41, 42]. Ritchie et al. in 2014 observed that the introduction of the small molecular ligand, 2−*{*[4 −(2− methylthiazol −4− ylmethyl) −[1, 4]diazepane −1 −carbonyl]amino} benzoic acid ethyl ester (*MTDB*), effectively decreased alternate folding by binding with the SARS-CoV −1 PRF pseudoknot [42]. Based on their experiment it was predicted that MTDB formed hydrogen bonds with the nucleotides in loop 3, inhibiting a non-native folding pathway. This binding resulted in reduction of the −1 PRF rate to near 0%. Recent in vitro experiments for SARS-CoV-2 determined that MTDB binding reduced −1 PRF by 60% [29]. This result was replicated and expanded upon by experiments that found various mutations in the SARS-CoV-2 −1 PRF pseudoknot sequence did not have a significant effect on anti-frameshifting activity of MTDB [27].

Mutations were reported in nearly every base in the SARS-CoV-2 −1 PRF pseudoknot sequences [27] available on GenBank [43] and GISAID [44]. A single nucleotide change from cytosine to uracil at position 62 (C62U) has been identified in an increasing proportion of sequences, up to 3% since the mutation was first recorded in March 2020 [26]. Due to the complicated nature of RNA folding in vivo, even a single nucleotide change can affect the resulting structure. Indeed, Neupane et al. identified a uracil to cytosine mutation at position 20 (U20C) for the SARS-CoV-2 −1 PRF pseudoknot that resulted in more than a three fold reduction in −1 PRF efficiency [27]. Therefore, mutations present in the population should be accounted for in structure prediction efforts for the SARS-CoV-2 −1 PRF pseudoknot, while currently they are not [25].

The mutations tested in the work of Neupane et al. [27], and which we further explore in this article, occur in regions that are important for pseudoknot structure. Three are located near the junction of stems (U20C, G29U, U58C), one is adjacent to an adenine bulge identified as critical in frameshifting [28] (C62U), and two are in Loop 2 where mutations have been shown previously to reduce frameshifting efficiency [45] (C43U, U47C). Incorporating known mutations in pseudoknot structure prediction will advance the understanding of potential conformations, which in turn could impact viral infectivity and pathology.

In addition to secondary structure, the 3-D conformation (e.g. triple-stranded helices) of RNA structures is critical to their function [46]. Existing *in silico* approaches to solve the three-dimensional RNA structure account for canonical and non-canonical base pairs, as well as pivotal tertiary interactions [47]. Many of the existing software packages to determine 3-D conformations of RNA also utilize secondary structure as an input constraint [30, 31, 32, 33, 34, 35]. Indeed, the latest 3-D physics-based modeling of the SARS-CoV-2 −1 PRF pseudoknot used the known native secondary structure as a starting point in identifying three unique stable conformations [25]. However, the non-native secondary structures of the SARS-CoV-2 −1 PRF pseudoknot have yet to be clearly identified and incorporated into structural prediction efforts. By presenting these predicted structures we provide alternate starting points and input constraints, which can improve the accuracy of 3-D physics-based modeling of the pseudoknot.

## 2 Materials and Methods

### 2.1 Data

The SARS-CoV reference genome NC_004718.3, SARS-CoV-2 reference genome NC_045512.2, and MERS-CoV reference genome NC_019843.3 were obtained from the National Center for Biotechnology Information (NCBI) [48].

### 2.2 Structural Similarity Method

RNAz [49], a de novo functional RNA detector, was used to predict structural similarity in an alignment of single-stranded RNA virus genomes that cause similar symptoms in patients [50, 51]. SARS-CoV-2, SARS-CoV, MERS-CoV, H1N1 (2009, ‘swine flu’), H5N1 (2006, Highly Pathogenic Avian Influenza), and two strains of Influenza A Virus (IAV) were included in the alignment.

The segmental Influenza A virus genomes were concatenated before alignment so that segments from the same virus would not align with any other segments from the same viral genome, and the reverse-complement was taken, as IAV is a negative-sense RNA virus. Concatenation locations were noted in order to later omit any predicted loci that spanned a concatenation point. These genomes were then aligned using the multiple sequence alignment tool Clustal Omega [52] version 1.2.4, with SARS-CoV-2 as the reference sequence.

The alignment was first pre-processed into overlapping windows using the rnazWindow.pl program. The output from rnazWindow.pl was used as input for RNAz, with the “–no-shuffle” option and the *P*-value cutoff set to 0.5. Note that the *P*-value used by RNAz is not a *P*-value in a statistical sense, but rather an ad hoc classification score in which the false positive rate was found to be *∼*4% for a cutoff of *P >* 0.5 and *∼*1% for a cutoff of *P >* 0.9 [53]. Output from RNAz was a file containing all windows with a positive RNAz signal and *P >* 0.5, as well as the predicted structures for each aligned sequence in the window and consensus structure. The accessory script rnazCluster.pl was then used on the RNAz output to group overlapping windows with predicted structure into loci. Finally, the output from rnazCluster.pl was used as input for rnazIndex.pl with the “–html” option, creating a “.html” file for visualization of the predicted loci.

### 2.3 Secondary Structure Prediction

Here we employ a pipeline (see Figure 4) with the goal of identifying the array of potential secondary structure conformations, both native and non-native, given an RNA sequence as input. We seek to leverage the hierarchical folding hypothesis by first predicting a set of energetically favourable initial stems for the input sequence (Figure 4, Steps 1 and 2) according to the energy parameter set by Andronescu et al. as implemented in HotKnots V2.0 [54, 55]. Based on the hierarchical folding view, these initial stems are expected to fold first, followed by additional stems [22]. The stems are given unique identifiers and sorted by free energy as calculated using the computeEnergy function from the HotKnots package [54, 55]. The initial stems are an intermediate output of the pipeline (see Figure 4). These predicted stems are then used as input constraints to guide the pseudoknotted secondary structure prediction (Figure 4, Step 3).

**Figure 4:**
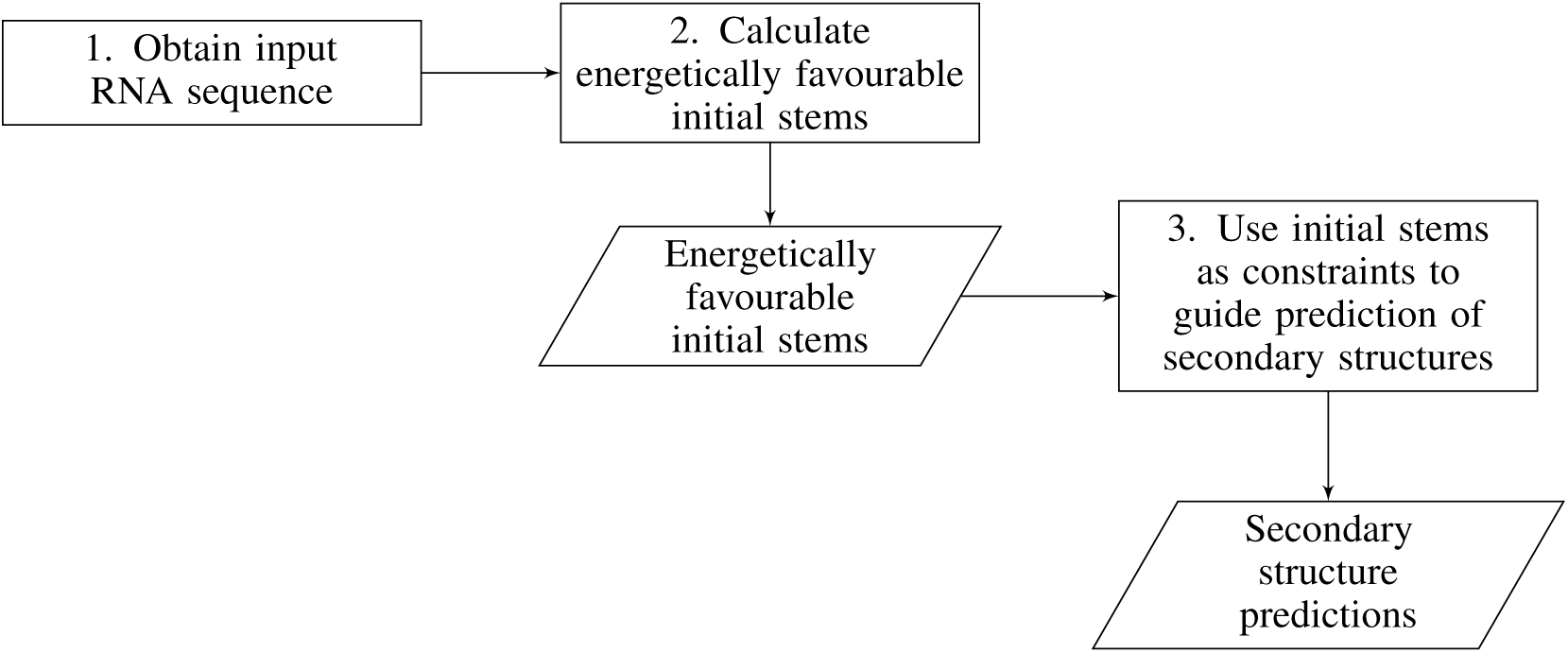
Hierarchical Folding Pipeline. Rectangles dictate actions, parallelograms denote an output.

While the initial stems have a significant impact in shaping the RNA structure, there is evidence that disrupting some initial base pairs can improve secondary structure prediction [56, 57]. Therefore, the Iterative HFold algorithm was selected to predict secondary structures (Figure 4, Step 3) [58]. This algorithm computes whether the input structure should be extended or disrupted based on predicting the minimum free energy structure. Iterative HFold is robust because it uses four different biologically sound methods to determine the structure with the lowest energy among all candidate structures. The value of employing multiple models instead of only a single method justifies the use of this specific algorithm.

Each initial candidate stem was used as an input structure for the Iterative HFold algorithm. The secondary structures produced based on each initial stem constraint are ranked by their minimum free energy. Each predicted secondary structure is linked to the initial stem(s) that produced it. We report initial stems and the resulting pseudoknotted structures for seven SARS-CoV-2 −1 PRF stimulating pseudoknot sequences: reference, and six mutated sequences (see Table 4). For the SARS-CoV-2 reference genome the 68 bp frameshifting pseudoknot sequence occurs at location 13475-13542 [29]. We also report initial stems and the resulting pseudoknotted structures for the MERS-CoV −1 PRF stimulating pseudoknot sequence. For the MERS-CoV reference genome the 71 bp frameshifting pseudoknot sequence occurs at location 13440-13510 [26].

## 3 Results

We first present the identified structural similarity between SARS-CoV, SARS-CoV-2, and MERS-CoV. We then enumerate the secondary structures of the SARS-CoV-2 −1 PRF pseudoknot for reference and mutated sequences. Finally we present the secondary structures predicted for the MERS-CoV −1 PRF pseudoknot.

### 3.1 Betacoronavirus −1 PRF Pseudoknot Structural Similarity

One predicted loci of structural similarity output from RNAz encompassed the region containing the −1 PRF pseudoknot for SARS-CoV-2. Structural similarity between SARS-CoV-2, SARS-CoV, and MERS-CoV was detected at the locations 13445 to 13557, 13375 to 13487, and 13410 to 13530 in their respective aligned genomes (see Table 1). The RNA-class probability as calculated by RNAz is 0.83, indicating that the probability this is a true alignment to be over 96%. The structural similarity locations for SARS-CoV-2 and MERS-CoV have a 100% overlap with their respective known −1 PRF pseudoknot sequences.

**Table 1:**
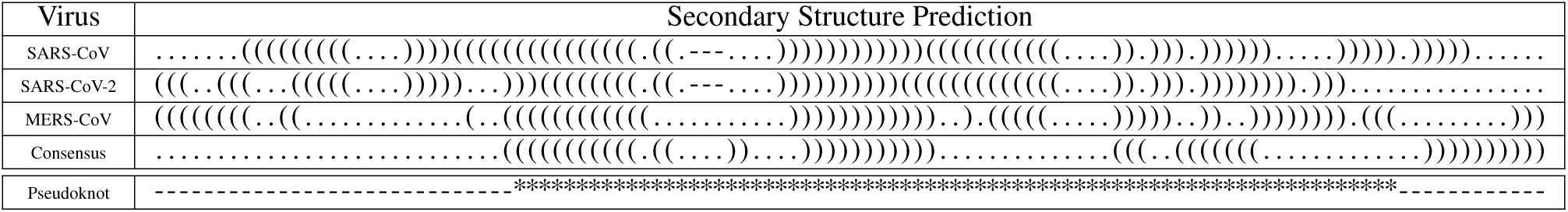
Structural similarity predictions for SARS-CoV, SARS-CoV-2, and MERS-CoV calculated using RNAz. Gaps in the alignment are represented as hyphen. Consensus structure is determined using an energy model augmented with covariance information. Asterisks (*) in bottom pseudoknot row correspond with known location of SARS-CoV-2 and MERS-CoV −1 PRF pseudoknot native structures.

### 3.2 SARS-CoV-2 −1 PRF Pseudoknot

Table 2 presents candidates of initial stems for the SARS-CoV-2 −1 PRF stimulating pseudoknot, as calculated based on the reference sequence. Using the 18 candidate initial stems as input constraints, 12 secondary structure predictions for the SARS-CoV-2 −1 PRF pseudoknot reference sequence are obtained using the Iterative HFold algorithm [58]. These predicted pseudoknotted structures are presented in Table 3 sorted by their free energies. In our predictions we use parentheses and brackets to demarcate pseudoknotted structures. Each period identifies an unpaired base. The first column in Table 3 lists the initial stem ID(s) corresponding with the constraint that resulted in the secondary structure prediction. Note that three different structures were each reached from more than one starting point, meaning the same structure resulted from different initial candidate stem constraints. The secondary structure with the lowest free energy as predicted by Iterative HFold is 100% consistent with the native structure.

**Table 2:**
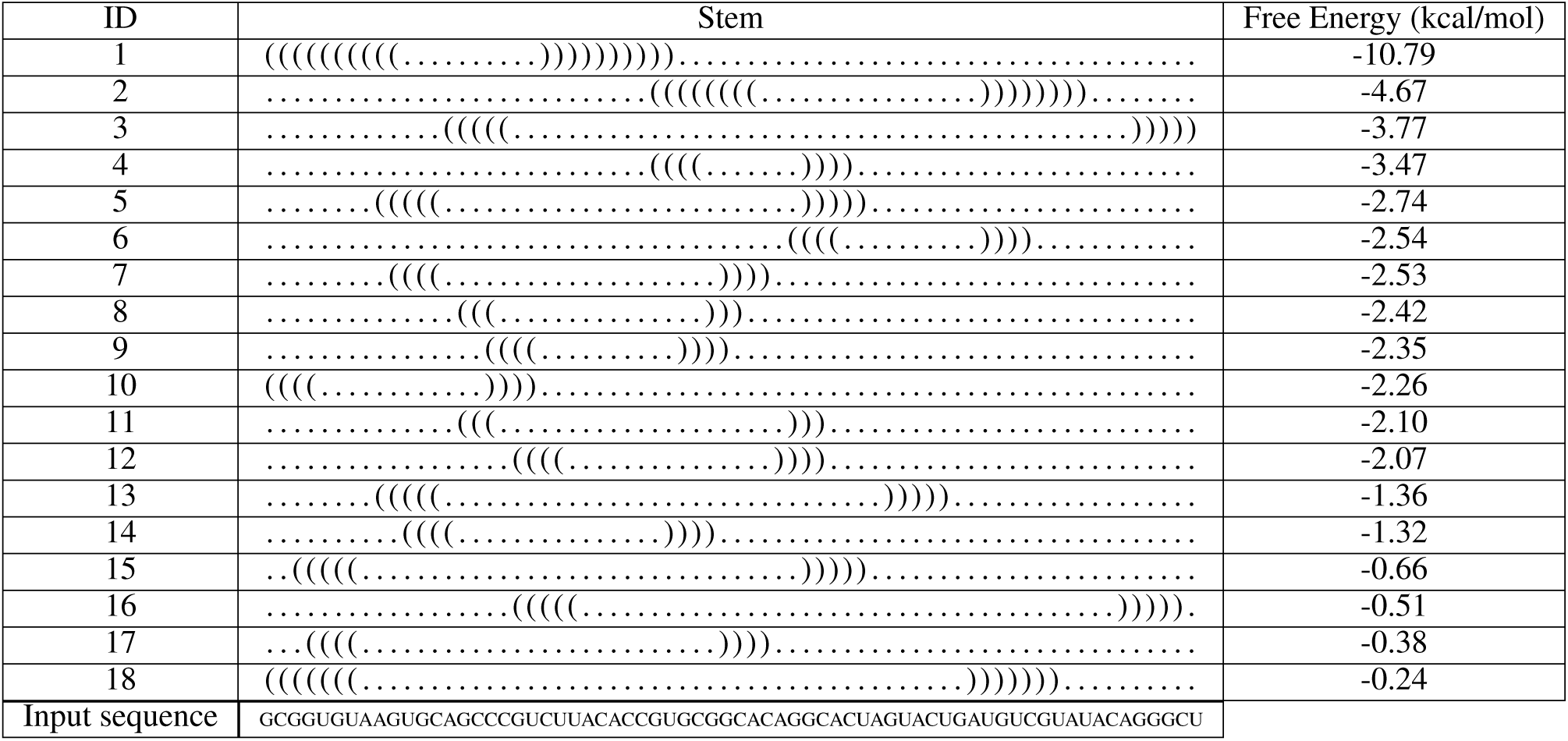
The predicted 18 most energetically favourable stems in SARS-CoV-2 −1 PRF stimulating pseudoknot reference sequence. These stems were used as structural guide for the Iterative HFold program to predict the SARS-CoV-2 secondary structure of the −1 PRF stimulating pseudoknot. First column provides stem ID and the third column provides free energy of the given stem. Free energies were calculated using the HotKnots V2.0 energy parameters of Andronescu et al. [55]. Input sequence provided in bottom row.

**Table 3:**
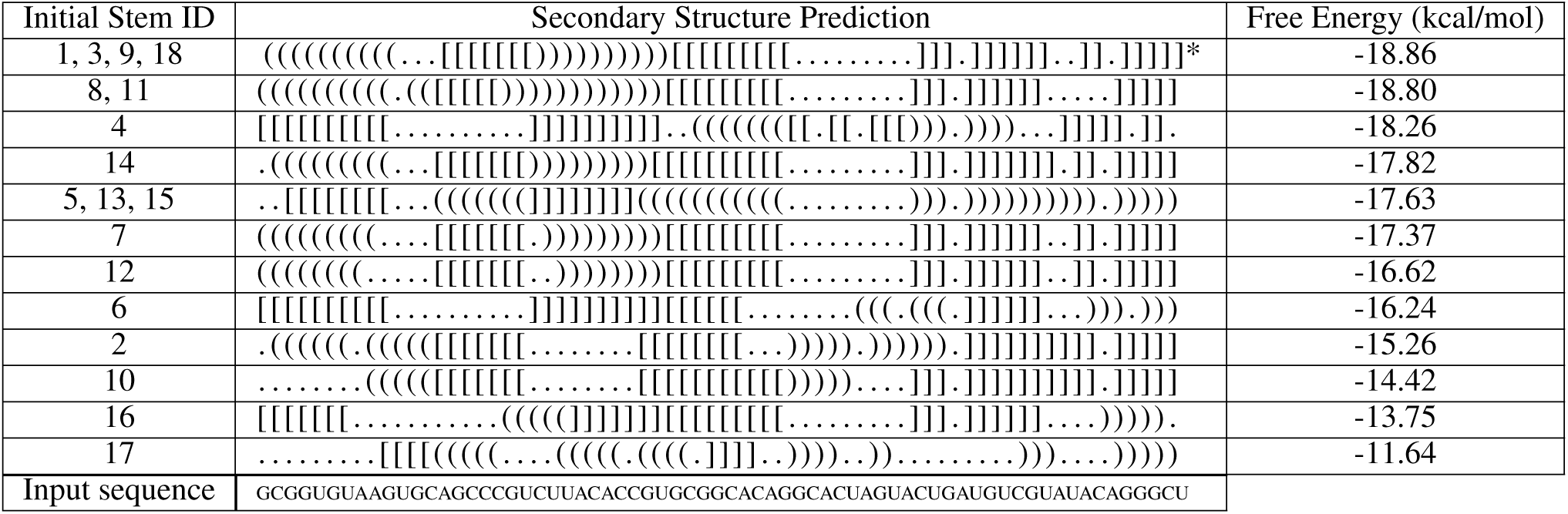
Predicted secondary structures for the SARS-CoV-2 −1 PRF stimulating pseudoknot based on the reference sequence. These structures are predicted by Iterative HFold given the input stems in Table 2 as structural constraints. Native structure is marked with an asterisk (*) in row 1. As shown in rows 1, 2 and 5 of the table, multiple initial stems can result in a single prediction for the −1 PRF stimulating pseudoknot. Input sequence provided in bottom row.

**Table 4:**
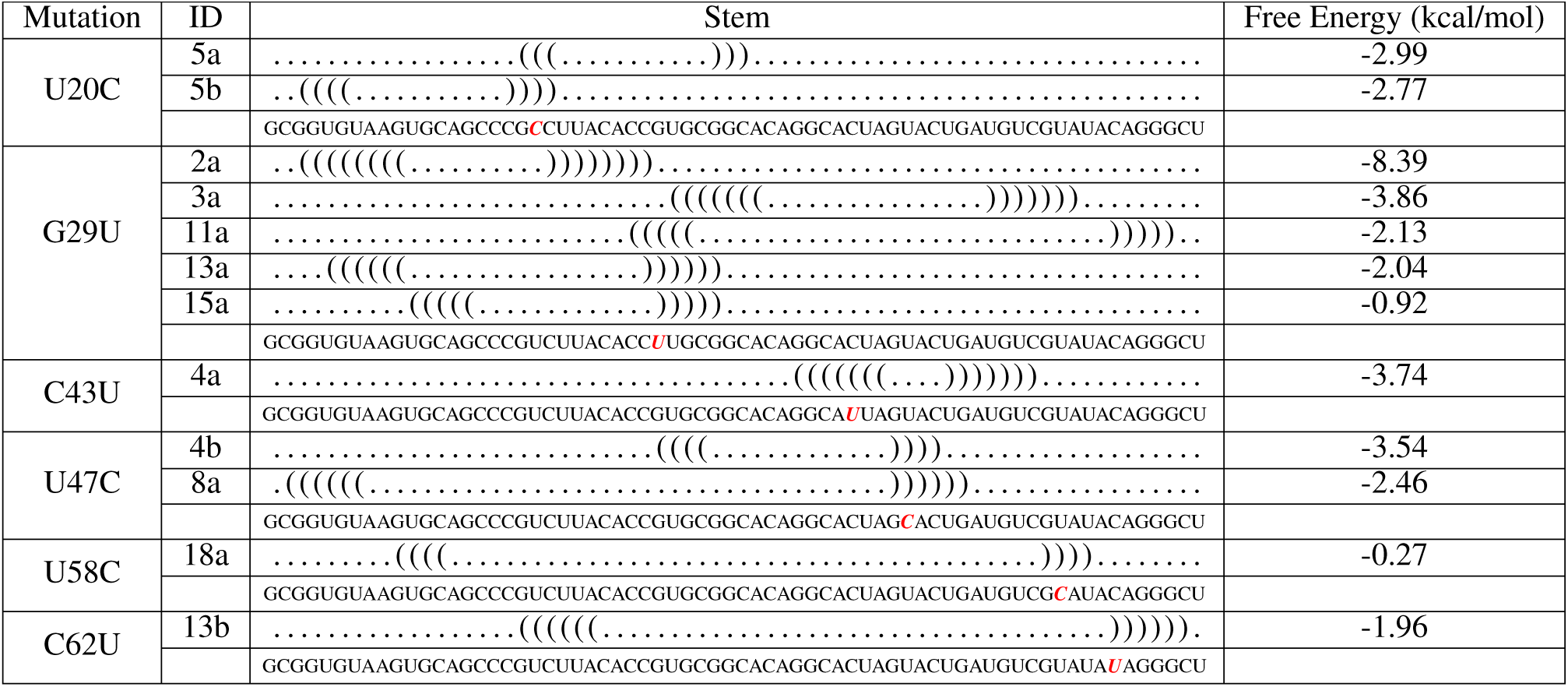
Predicted most energetically favourable stems for SARS-CoV-2 −1 PRF stimulating pseudoknot mutated sequences. These stems were used as a structural guide for the Iterative HFold program to predict the SARS-CoV-2 secondary structure of the −1 PRF stimulating pseudoknot. Predictions in each row are based on the sequence with the mutation given in the first column. Second column provides stem ID and the fourth column provides free energy of the given stem. Stem IDs here correspond with stem free energy relative to the reference sequence initial stems. For example, in row one the stem has a free energy of −2.99 and is denoted by stem ID 5*a*. Relative to the stems predicted for the reference sequence (see Table 2), this stem has the fifth lowest free energy. Free energies calculated using the HotKnots V2.0 energy parameters of Andronescu et al. [55]. Input sequence provided in bottom row of each mutation section, respective mutations italicized in red.

We repeated the same steps with the mutated sequences. The predicted initial stems did change in some cases between the reference sequence and the mutated sequences. Certain stems that were predicted for both the reference sequence and respective mutated sequences had a different calculated free energy based on the mutated sequence. Some stems that were identified as energetically probable for the reference sequence were not found as energetically probable for specific mutated sequence. In addition, some mutated sequences led to initial stem predictions that were not present for the reference sequence. We present initial stems for mutated sequences with an ID based on where they would have been ranked by free energy relative to the reference sequence stems (see Table 4).

Initial stem predictions for the U20C mutated sequence changed the predicted energy of five different stems. Stem 1 free energy increased from −10.79 to −10.28. The predicted energies for stems 9, 10, 12, and 16 all decreased. Stem 9 decreased from −2.35 to −3.87, stem 10 decreased from −2.26 to −3.40, stem 12 decreased from −2.07 to −3.85, and stem 16 decreased from −0.51 to −2.29. Two novel stems were detected referred to as 5*a* and 5*b* in Table 4.

Repeating the method for the sequence with the guanine/uracil mutation at position 29 (G29U) did not change the predicted energy of any stems, but did result in four previously detected stems that were not detected (1, 2, 4 and 14). Five novel stems were detected based on the sequence with the G29U mutation, referred to as 2*a*, 3*a*, 11*a*, 13*a*, and 15*a* in Table 4.

For the sequence with the cytosine/uracil mutation at position 43 (C43U), two stems changed in their predicted energy. Stem 4 increased in free energy from −3.47 to −1.62, as did stem 5, from −2.74 to −0.91. Stems 6 and 15 were not detected for C43U, and a novel stem was detected, referred to as 4*a* in Table 4.

Stem predictions for the sequence with the uracil/cytosine mutation at position 47 (U47C) decreased the predicted energy of stem 13 from −1.36 to −3.32. Two novel stems were detected, referred to as 4*b* and 8*a* in Table 4.

For the sequence with the uracil/cytosine mutation at position 58 (U58C), two stems decreased in energy: stem 2 decreased from −4.67 to −6.63 and stem 18 decreased from −0.24 to −1.88. One novel stem was detected, referred to as 18*a* in Table 4.

Finally, for the sequence with the cytosine/uracil mutation at position 62 (C62U), stem 16 was not detected. There was a novel stem identified referred to as 13*b* in Table 4.

We used the 18 candidate stems from the reference sequence (see Table 2) in addition to novel candidate stems for respective mutated sequences (see Table 4) as input constraints to predict secondary structures using the Iterative HFold algorithm [58]. For each of the six SARS-CoV-2 −1 PRF pseudoknot mutated sequences, either 12 or 13 secondary structures were predicted. However, the predicted secondary structures are markedly different from those predicted based on the reference sequence. The native structure, the structure with lowest free energy in Table 3, was not predicted for U20C, G29U, or C62U mutated sequences based on any of the initial stems. For example, stems 1 and 3 that result in the native structure based on the reference sequence, did not result in the native structure based on the sequence with the U20C mutation (see Table 5). Instead, stems 1 and 3 now join stems 4, 11 and 16 resulting in a non-native structure. Due to the U20C mutation, Stems 9 and 18 cannot achieve the native structure, and each are predicted to lead to new non-native structures. Three structure clusters (identical prediction from multiple different initial stems) were identified for each mutated sequence, except for the U58C mutation that led to five predicted structure clusters. As expected, predictions based on Loop 2 mutations C43U and U47C were similar.

**Table 5:**
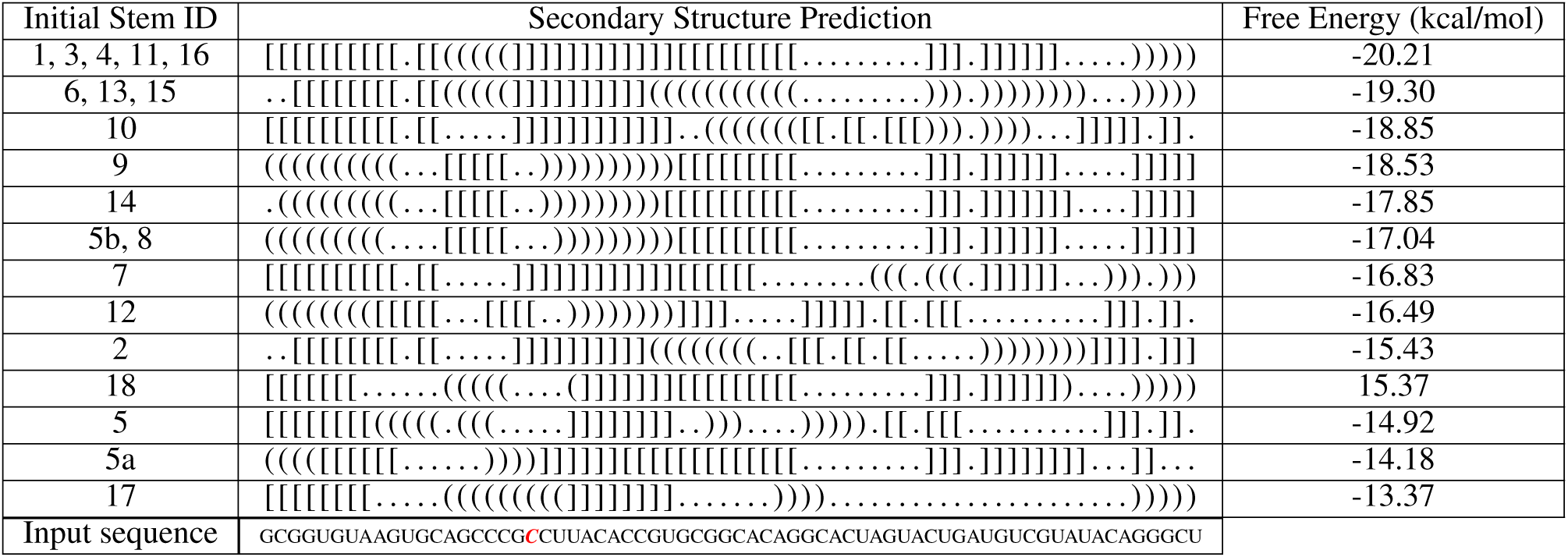
Predicted secondary structures for SARS-CoV-2 −1 PRF stimulating pseudoknot with U20C mutation. These structures are predicted by Iterative HFold given the input structures in Table 2, as well as stems 5*a* and 5*b* in Table 4, as structural constraint. Input sequence provided in bottom row, C mutation italicized in red.

**Table 6:**
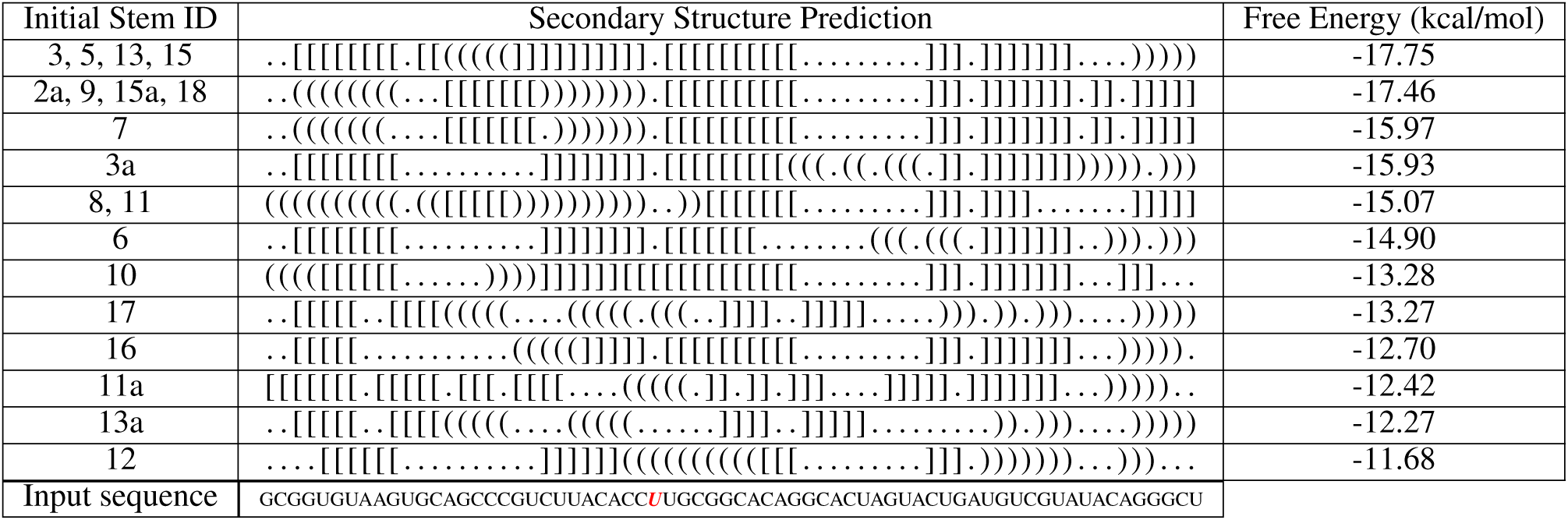
Predicted secondary structures for SARS-CoV-2 −1 PRF stimulating pseudoknot with G29U mutation. These structures are predicted by Iterative HFold given the input structures in Table 2 (stems 1, 2, 4, and 14 omitted), as well as stems 2*a*, 3*a*, 11*a*, 13*a* and 15*a* in Table 4, as structural constraint. Input sequence provided in bottom row, U mutation italicized in red.

**Table 7:**
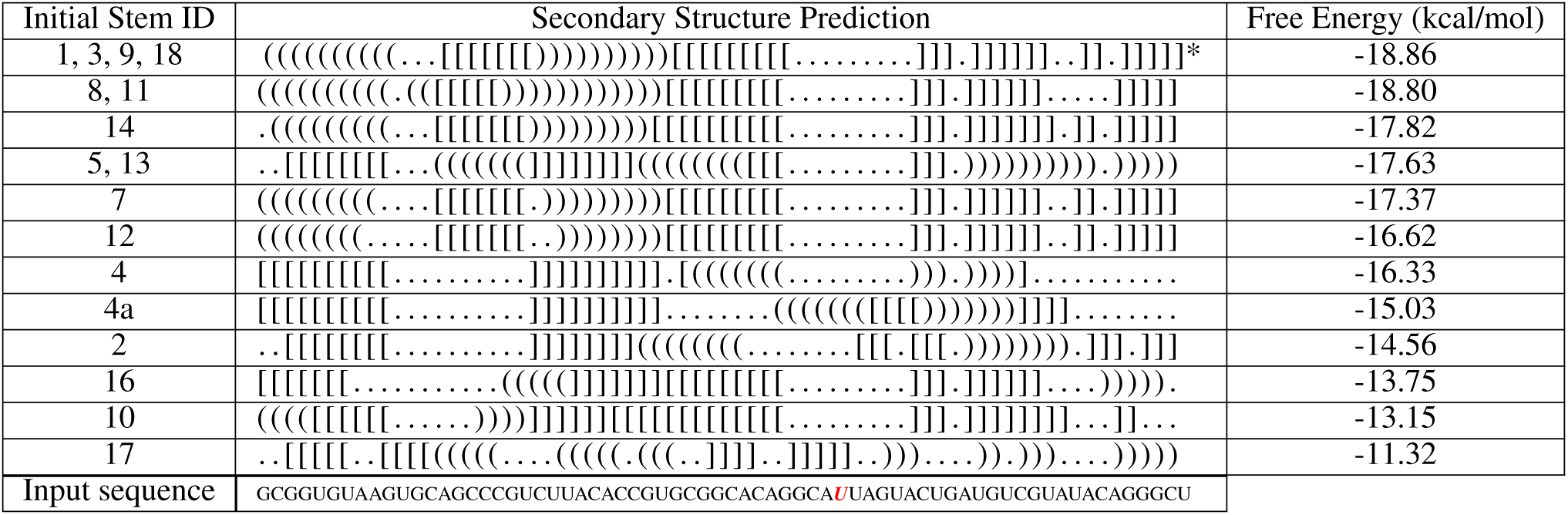
Predicted secondary structures for SARS-CoV-2 −1 PRF stimulating pseudoknot with C43U mutation. These structures are predicted by Iterative HFold given the input structures in Table 2 (stems 6 and 15 omitted), as well as stem 4*a* in Table 4, as structural constraint. Native structure is marked with an asterisk (*) in row 1. Input sequence provided in bottom row, U mutation italicized in red.

**Table 8:**
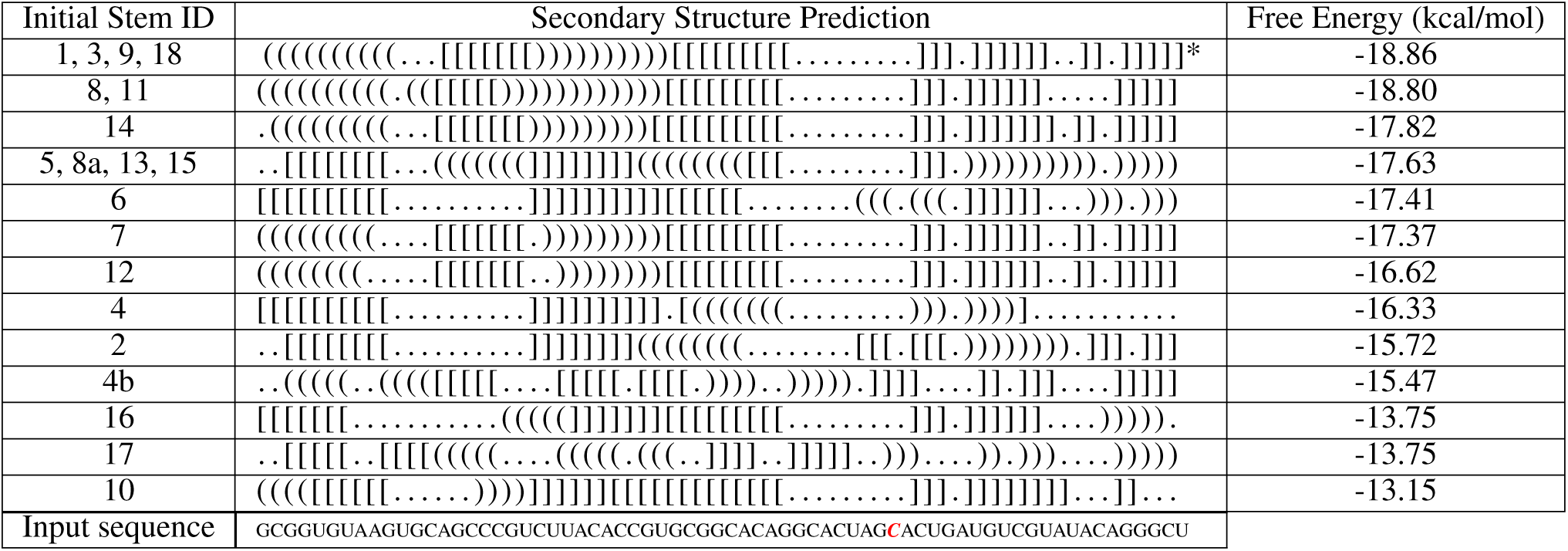
Predicted secondary structures for SARS-CoV-2 −1 PRF stimulating pseudoknot with U47C mutation. These structures are predicted by Iterative HFold given the input structures in Table 2, as well as stem 4*b* and 8*a* in Table 4, as structural constraint. Native structure is marked with an asterisk (*) in row 1. Input sequence provided in bottom row, C mutation italicized in red.

**Table 9:**
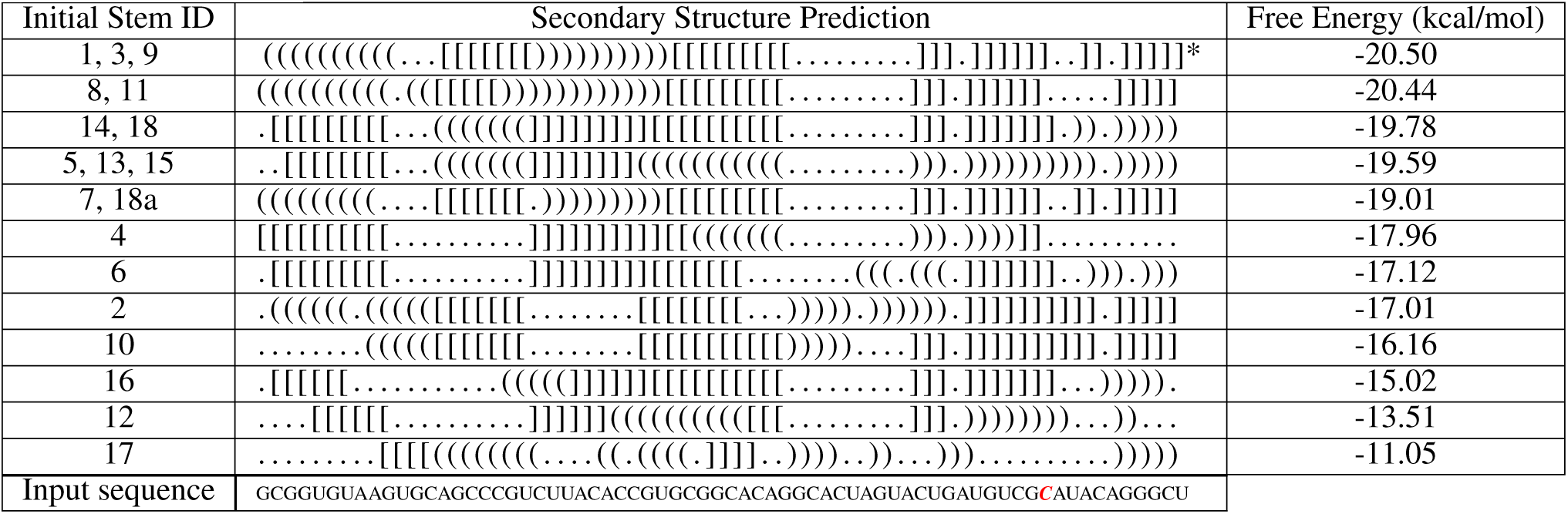
Predicted secondary structures for SARS-CoV-2 −1 PRF stimulating pseudoknot with U58C mutation. These structures are predicted by Iterative HFold given the input structures in Table 2, as well as stem 18*a* in Table 4, as structural constraint. Native structure is marked with an asterisk (*) in row 1. Input sequence provided in bottom row, C mutation italicized in red.

**Table 10:**
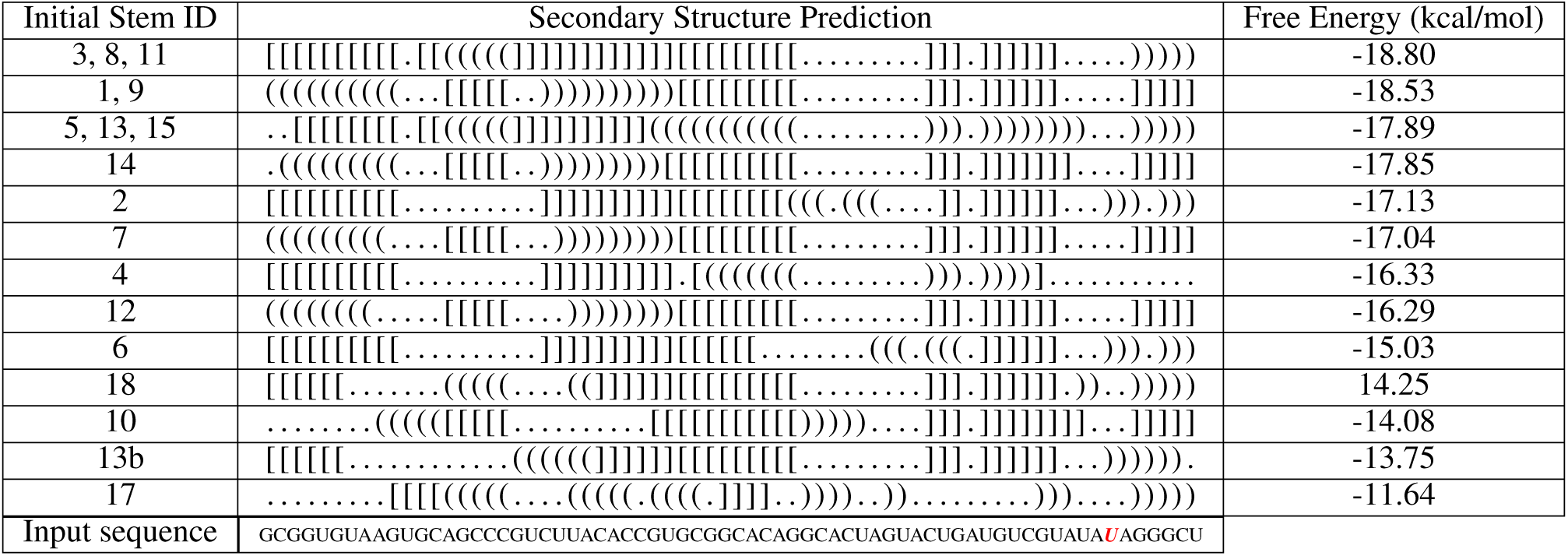
Predicted secondary structures for SARS-CoV-2 −1 PRF stimulating pseudoknot with C62U mutation. These structures are predicted by Iterative HFold given the input structures in Table 2 (stem 16 omitted), as well as stem 13*b* in Table 4, as structural constraint. Input sequence provided in bottom row, U mutation italicized in red.

### 3.3 MERS-CoV-2 −1 PRF Pseudoknot

Table 11 presents candidate initial stems for the MERS-CoV-2 −1 PRF stimulating pseudoknot, as calculated based on the reference sequence. Using the 19 candidate initial stems (see Table 11) as input constraints, 13 secondary structure predictions for the MERS-CoV-2 −1 PRF pseudoknot reference sequence are obtained using the Iterative HFold algorithm [58]. These predicted pseudoknotted structures are presented in Table 12 sorted by free energy. The first column in Table 12 indicates the initial stem ID(s) corresponding with the constraint that resulted in the secondary structure prediction. Note that four structures were reached from more than one starting point (see rows 1, 2, 6, and 7 in Table 12). There is no well established native secondary structure for the MERS-CoV −1 PRF pseudoknot. The secondary structure with the lowest free energy as predicted by Iterative HFold is 93% consistent with the structure presented by Fourmy and Yoshizawa [26].

**Table 11:**
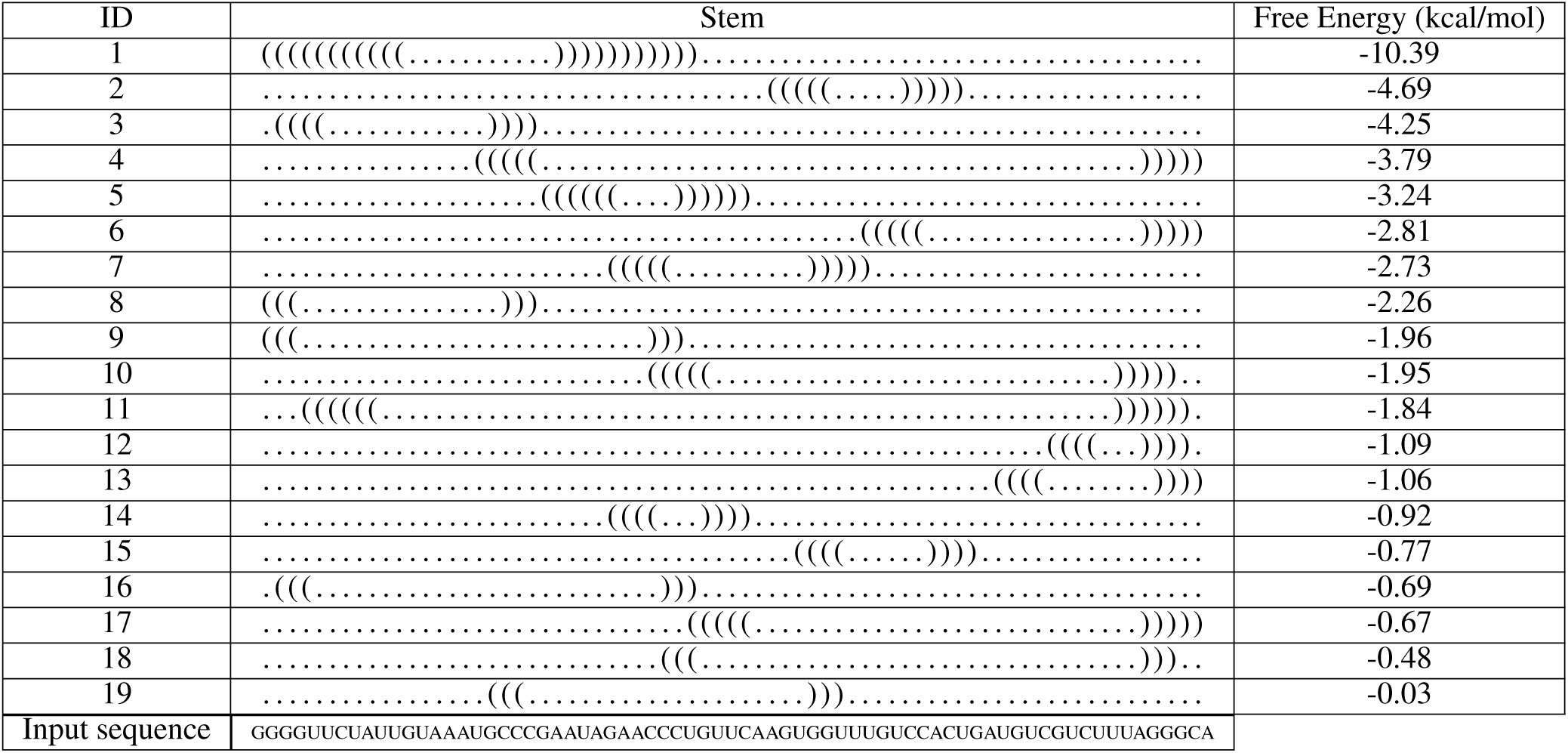
Predicted most energetically favourable stems for MERS-CoV −1 PRF stimulating pseudoknot reference sequence. These stems were used as a structural guide for the Iterative HFold program to predict the secondary structure of the −1 PRF stimulating pseudoknot. First column provides stem ID and the third column provides free energy of the given stem. Free energies calculated using the HotKnots V2.0 energy parameters of Andronescu et al. [55]. Input sequence provided in bottom row.

**Table 12:**
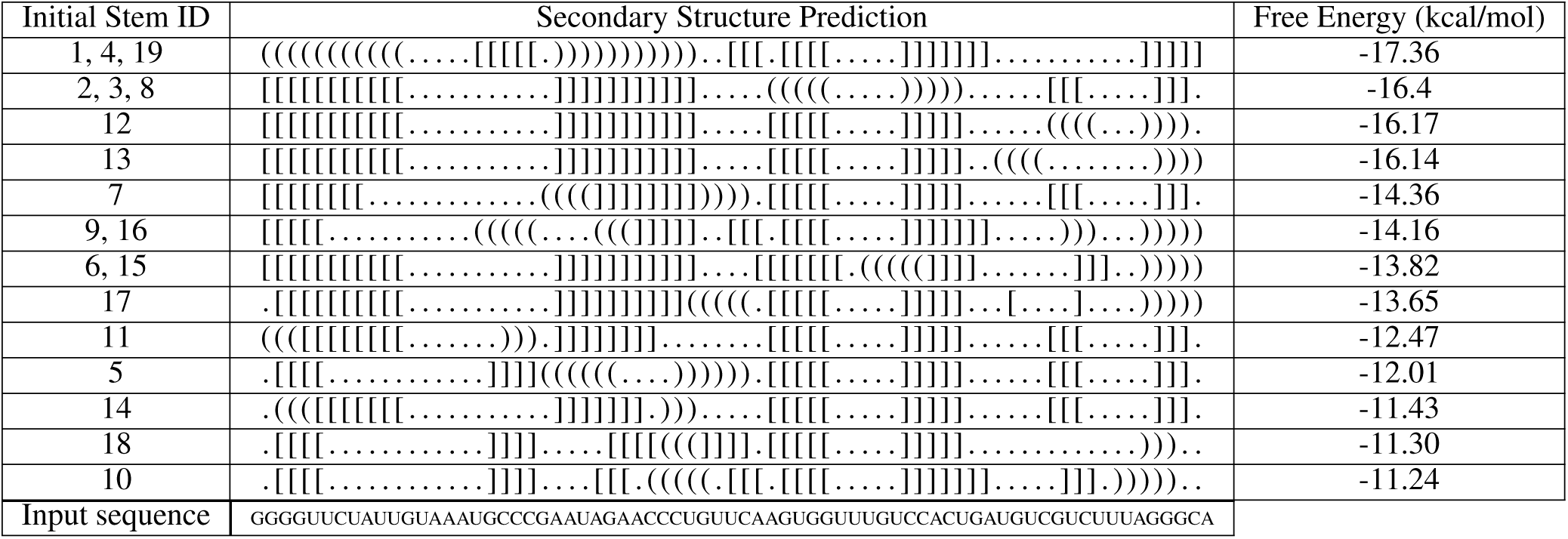
Predicted secondary structures for MERS-CoV −1 PRF stimulating pseudoknot based on the reference sequence. These structures are predicted by Iterative HFold given the input stems in Table 11 as structural constraints. Input sequence provided in bottom row.

## 4 Discussion

We set out to detect structural similarity between different coronaviruses. Here we aligned viral genomes and used RNAz to identify a region of structural similarity for three Betacoronaviruses: SARS-CoV, SARS-CoV-2, and MERS-CoV. It is expected that SARS-CoV and SARS-CoV-2 have the structural similarity we detected in the area of the −1 PRF pseudoknot, because the pseudoknot sequence is 99% identical between the two. In addition to similarity between SARS-CoV and SARS-CoV-2, we detected structural similarity with MERS-CoV in its −1 PRF pseudoknot region. While this result was recently hypothesized by Fourmy and Yoshizawa [26], to the best of our knowledge so far it has not been experimentally validated. In this work, we add more evidence for this structural similarity based on a replicable method. Furthermore, the RNAz output perfectly aligned the 68 and 71 bp −1 PRF pseudoknot sequences for SARS-CoV-2 and MERS-CoV. This is interesting because there is no sequence similarity between the SARS-CoV-2 and MERS-CoV −1 PRF sequences as detected using the Basic Local Alignment Search Tool [59]. Similarities in structure between coronaviruses may contain vital information that can validate proposed explanations for frameshifting. The individual and consensus structures presented here can further inform structure prediction, providing previously unavailable insights to explain pathways for structure formation.

Beyond structural similarity, we identify the array of potential secondary structure conformations, both native and non-native, for the SARS-CoV-2 and MERS-CoV −1 PRF pseudoknots. Here we employed a hierarchical folding based method for prediction of the secondary structures. We first identified energetically favourable initial stems for a given RNA sequence. We then used the initial stems to guide our secondary structure predictions. We presented pseudoknot structural predictions based on the reference sequence of the SARS-CoV-2 −1 PRF pseudoknot, as well as six mutated sequences. These mutated sequences were recently tested for their effect on −1 PRF efficiency [27].

Our results show that, while the native structure of the pseudoknot has the lowest free energy, there are multiple predicted non-native structures that are comparable both from an energy and structural standpoint. This demonstrates the flexibility and conservation of the three-stemmed structure. Specific to the SARS-CoV-2 −1 PRF pseudoknot, further investigation is needed to determine the frequency that different non-native secondary structures are realized. Focus should be placed on understanding the refolding dynamics of partially unfolded structures, which may mimic the interplay of the ribosome and pseudoknot in −1 PRF.

In addition to non-native structures that are similar to the native structure, we predicted non-native structures that are very different from the native expectation. These non-native secondary structures are energetically close to the native structure and likely play a role in regulating frameshifting. In both SARS-CoV-2 and MERS-CoV, the second most energetically favourable stem (Stem ID 2, see Tables 2 and 11) results in a non-native structure prediction that is very different than the native expectation (see Table 3 and 12). Although stem 2 is not the most energetically probable, its formation is supported by our structural alignment. While others discussed non-native structures and their role in frameshifting [23], those structures have yet to be explicitly identified for further validation. Further experiments are needed to confirm the exact structure of non-native conformations for the SARS-CoV-2 −1 PRF pseudoknot and how these specific structures correlate with frameshifting efficiency.

For SARS-CoV-2 reference sequence predictions, three predicted pseudoknotted structures including the native structure, resulted from multiple initial stem input constraints. These structures can be accessed from different starting points, which may indicate an increased likelihood for these specific pathways. Our identification of possible pathways by which each pseudoknotted structure is formed advances existing understanding of structure formation.

Secondary structure predictions based on the SARS-CoV-2 −1 PRF pseudoknot mutated sequences are demonstrably different than those for the reference sequence. Depending on the location of the mutation, different novel secondary structures were predicted. In some cases, initial stems that resulted in one structure predicted for the reference sequence, resulted in a different structure predicted based on the mutated sequence. Mutations also changed the free energy for predicted structures that were consistent between reference and mutated sequence. Interestingly, while there were changes in predicted secondary structures, the function of the pseudoknot is expected to be conserved for all mutations other than U20C [27]. Given that the structure of the pseudoknot with sequence mutations is different but still functional, we hypothesize that the relationship between pseudoknot structure and function may be more dynamic and flexible than previously expected.

In the case of the U20C mutation, we identify a structure that is very different from the native expectation and has a low free energy (Table 5 row 3, structure predicted from stem 10 with free energy −18.85). This highly non-native structure is ranked third in terms of free energy, which is not the case for any other mutation. Interestingly, U20C was the only mutation that has been determined to result in significant decrease in −1 PRF efficiency [27]. Further experiments are needed to validate this result and confirm the structural cause for the decrease in −1 PRF efficiency based on the U20C mutation. It is important to investigate exactly which conformations of the pseudoknot are responsible for inducing frameshifting. As existing and new SARS-CoV-2 mutations become more prevalent, determining their impact on −1 PRF pseudoknot structures is key to long-term success of small molecule therapy.

Finally, RNA tertiary contact prediction tools are poised to play an essential role in developing and vetting small molecule therapies. Indeed, the most recent efforts in identifying unique stable 3D conformations of the SARS-CoV-2 −1 PRF pseudoknot can have a great impact in treatment development [25]. We believe such efforts can benefit from a more comprehensive overview of the SARS-CoV-2 1 PRF pseudoknot secondary structure landscape, one that includes non-native structures. There are many RNA tertiary contact prediction tools that are equipped to incorporate knowledge of secondary structure as simulation constraints [30, 31, 32, 33, 34, 35]. Given the complicated nature of modeling tertiary interactions, the initialization of such simulations should account for multiple initial secondary structures of the pseudoknot. Our results provide pseudoknot-free constraints (Tables 2 and 4), as well as pseudoknotted constraints (Tables 3, 5-10) that can be used as starting points or to provide context to future SARS-CoV-2 structure prediction efforts.

## 5 Conclusion

We identified structural similarity related to the −1 PRF pseudoknots of SARS-CoV, SARS-CoV-2, and MERS-CoV. Our structural alignment confirms the connection between the SARS-CoV-2 and MERS-CoV −1 PRF pseudoknots. We followed the hierarchical folding hypothesis to explore the structural landscape of the SARS-CoV-2 and MERS-CoV −1 PRF pseudoknots. Here, we first identified energetically favourable initial stems for each sequence, then incorporated the initial stem information into secondary structure predictions for the SARS-CoV-2 and MERS-CoV −1 PRF pseudoknots. Previous work to model the SARS-CoV-2 −1 PRF pseudoknot has acknowledged that non-native structures are relevant [23], but this concept has not been fully integrated into the latest prediction efforts [25]. In this work, we identified non-native pseudoknotted structures, as well as possible pathways by which each predicted structure is realized. This holistic methodology may advance the understanding of the SARS-CoV-2 −1 PRF pseudoknot.

We utilized knowledge of mutations in the SARS-CoV-2 −1 PRF pseudoknot sequence [26] to contrast structure prediction between the reference and mutated sequences. Small secondary structure differences can have a profound impact on resulting tertiary interactions and three-dimensional conformations of the pseudoknot. Our structural predictions can be further utilized in 3-D physics-based modeling of pseudoknots as constraints [30, 31, 32, 33, 34, 35].

## 6 Author Contributions

LT and HJ designed the research. LT and LL carried out all simulations and modeling. LT was the primary article writer. All authors contributed to the revision process.

## 7 Acknowledgments

We thank and acknowledge the University of Victoria (UVic), Department of Computer Science for sponsoring this research through the UVic Fellowship Award. We also thank the Computational Biology Research and Analytics Lab for invaluable feedback.

## 8 Supplementary Material

An online supplement to this article can be found by visiting our github repository at https://github.com/HosnaJabbari/frameshifting.

## Notes

https://github.com/HosnaJabbari/frameshifting

### Competing Interest Statement

The authors have declared no competing interest.

